# Evolving in the darkness: phylogenomics of *Sinocyclocheilus* cavefishes highlights recent diversification and cryptic diversity

**DOI:** 10.1101/2021.10.09.460971

**Authors:** Tingru Mao, Yewei Liu, Mariana M. Vasconcellos, Marcio R. Pie, Gajaba Ellepola, Chenghai Fu, Jian Yang, Madhava Meegaskumbura

## Abstract

Troglomorphism— any morphological adaptation enabling life to the constant darkness of caves, such as loss of pigment, reduced eyesight or blindness, over-developed tactile and olfactory organs—has long intrigued biologists. However, inferring the proximate and ultimate mechanisms driving the evolution of troglomorphism in freshwater fish requires a sound understanding of the evolutionary relationships between surface and troglomorphic lineages. We use Restriction Site Associated DNA Sequencing (RADseq) to better understand the evolution of the *Sinocyclocheilus* fishes of China. With a remarkable array of derived troglomorphic traits, they comprise the largest cavefish diversification in the world, emerging as a multi-species model system to study evolutionary novelty. We sequenced a total of 120 individuals throughout the *Sinocyclocheilus* distribution. The data comprised a total of 646,497⍰bp per individual, including 4378 loci and 67,983 SNPs shared across a minimum of 114 individuals at a given locus. Phylogenetic analyses using either the concatenated RAD loci (RAxML) or the SNPs under a coalescent model (SVDquartets, SNAPP) showed a high degree of congruence with similar topologies and high node support (> 95 for most nodes in the phylogeny). The major clades recovered conform to a pattern previously established using Sanger-based mt-DNA sequences, with a few notable exceptions. We now recognize six major clades in this group, elevating the blind cavefish *S. tianlinensis* and the micro-eyed *S. microphthalmus* as two new distinct clades due to their deep divergence from other clades. PCA plots of the SNP data also supports the recognition of six major clusters of species congruent with the identified clades based on the spatial arrangement and overlap of the species in the PC space. A Bayes factor delimitation (BFD) analysis showed support for 21 species, recognizing 19 previously described species and two putative new cryptic ones. Two species whose identities were previously disputed, *S. furcodorsalis* and *S. tianeensis*, are supported here as distinct species. In addition, our multi-species calibrated tree in SNAPP suggests that the genus *Sinocyclocheilus* originated around 10.5 Mya, with most speciation events occurring in the last 2 Mya, likely favored by the uplift of the Qinghai-Tibetan Plateau and cave occupation induced by climate-driven aridification during this period. These results provide a firm basis for future comparative studies on the evolution of *Sinocyclocheilus* and its adaptations to cave life.

## 1. INTRODUCTION

Many animal lineages across the world, ranging from flatworms (Leal-Zanchet et al., 2014) to fish and amphibians (Hutchison, 1958), have evolved to live in caves. In order to occupy cave subterranean habitats, each of these lineages had to adapt to extreme conditions of low availability of food, oxygen, and light, leading to the repeated acquisition of one or more adaptations, including elongated appendages, lowered metabolism, specialized sensory systems, and loss of eyes and pigmentation (Jeffery, 2019). The concerted evolution of these traits across different taxa, giving rise to convergent troglomorphic forms, comprise a great example of the process of natural selection in response to cave-associated selective regimes (Borowsky, 2010; Jeffery et al., 2010; Klaus et al., 2013; Porter et al., 2003). Though large troglomorphic radiations are rare – mainly because of the limited extent of cave habitats – the cyprinid fish genus *Sinocyclocheilus* shows an exceptional diversity of cave adaptations including at least three independent origins of cave-adapted phenotypes (Mao et al., 2021).

With 75 known species, *Sinocyclocheilus* includes the largest radiation of cavefish in the world, which diversified across Karstic habitats associated with the Li River in the Guangxi, Guizhou, and Yunnan provinces of China (Jiang et al., 2019; Zhao and Zhang, 2009). The troglomorphic adaptations of *Sinocyclocheilus* include the development of a specialized sensory system, such as degeneration or complete loss of eyes, degeneration of scales, loss of pigmentation, expansion of pectoral fins, evolution of ‘horns’ and enhancement of the neuromast system (He et al., 2013; Li et al., 2020; Ma et al., 2020; Meng et al., 2013). This wealth of troglomorphic adaptations makes *Sinocyclocheilus* an important multi-species model system to study evolutionary novelty in response to selection (Chen et al., 2009; Meng et al., 2013; Xiao et al., 2005; Yang et al., 2016). Yet, this requires a robust time-calibrated phylogeny for this group including several independent markers across their genome, which is currently missing.

Our current understanding of the phylogenetic relationships within *Sinocyclocheilus*, based solely on mitochondrial DNA (mtDNA), revolves around the recognition of four main clades, A, B, C, and D (Mao et al., 2021), sometimes also referred to as the “jii”, “angularis”, “cyphotergous”, and” tingi” groups (Li and He, 2009; Xiao et al., 2005; Zhao and Zhang, 2009). These are sequential clades rather than reciprocally monophyletic units. The earliest emerging clade (Clade A) is restricted to Guangxi, at the eastern fringes of the genus distribution. Clades B and C have overlapping distributions restricted to the middle of the genus distribution, and species of Clade D are found mostly in the lotic habitats associated with hills to the west. In addition, Clade B encompasses most species with extensive troglomorphic traits such as the complete loss of eyes (blind) and well-formed forehead protrusions (horns). The genus is thought to have originated during the Miocene-Pliocene with cave occupation taking place predominantly during the Pliocene-Pleistocene transition in response to an uplift event and aridification of the Qinghai-Tibetan Plateau (Mao et al., 2021).

Nearly all of the molecular studies to date on the phylogenetic relationships within *Sinocyclocheilus* have been based on mtDNA (Chen et al., 2018; Jiang et al., 2019; Liang et al., 2011), limiting some of the diversification-scale analyses as well as insights into the population-level processes that shape this remarkable radiation. This is an important limitation, given that the uniparental nature of mtDNA inheritance might obscure important events of admixture in the history of *Sinocyclocheilus* while hampering the inference of species limits given that it represents only a single genetic locus. With the advent of next-generation sequencing, particularly highly efficient methods for recovering thousands of orthologous loci such as Restriction-site associated DNA sequencing (Cariou et al., 2013), we can infer the relationships within *Sinocyclocheilus* with higher accuracy, as well as its species limits, ancestral admixture, and diversification events.

The main goals of our study were: (1) to infer the evolutionary relationships and divergence times within *Sinocyclocheilus* through phylogenomic methods; (2) to assess the level of genetic support for the currently recognized species in the genus using Bayesian species delimitation methods; and (3) to investigate possible ancestral admixture events among *Sinocyclocheilus* species. In short, we were able to build a well-resolved tree for the genus despite recognizing some introgression events, to confirm the distinctiveness of 19 previously described species, and to recognize two additional cryptic species. Finally, we estimate that the most recent common ancestor (MRCA) of *Sinocyclocheilus* originated around 10.5 Mya, which is considerably older than previous estimates based on mitochondrial data alone.

## 2. METHODS

### 2.1 Taxon sampling and laboratory work

To elucidate the phylogenetic relationships of *Sinocyclocheilus* using multiple nuclear genomic markers, we carried out a detailed phylogenomic analysis of the group .We collected samples of 120 individuals in Guangxi, Yunnan and Guizhou provinces of China, including 19 previously recognized species and 2 unidentified species of *Sinocyclocheilus* (Fig. S1). Given the rarity of these species, only fin-clips were collected from all individuals and frozen immediately at −85 °C until DNA extraction. Genomic DNA was extracted using the DNeasy Blood and Tissue Kit (Qiagen Inc., Valencia, CA) following the manufacturer’s protocols. Electrophoresis was performed to ensure DNA integrity of the samples prior to genomic library preparation.

We employed a RAD sequencing protocol (Baird et al., 2008), which involves the use of a single restriction enzyme to cut genomic DNA at specific sites, using molecular barcodes to identify each individual and build a genomic library of short fragments distributed across the entire genome of all individuals. Each DNA sample was digested using the EcoRI enzyme and ligated to a P1 Illumina adapter at the compatible ends of the fragments. This adapter allows forward amplification with Illumina sequencing primers containing a 6-bp long nucleotide barcode for sample identification. The barcoded adapter-ligated fragments from all individuals were subsequently pooled, randomly sheared, and size-selected. DNA was then ligated to a second P2 adapter, a Y-shaped adapter with divergent ends. Finally, fragment sizes of 200-400 bp and 400-600 bp were isolated for library construction using a MinElute Gel Extraction kit (Qiagen). We quantified DNA concentration in a Qubit2.0, and checked the quality (insert size) of genomic libraries in an Agilent 2100. qPCR was also performed to detect the effective concentration of libraries (if > 2nM) in the appropriate insert size. Final genomic libraries were then sequenced on an Illumina HiSeq2500 platform generating 125-bp paired-end reads.

### 2.2 RADseq data assembly

Our pre-processing bioinformatics of the sequenced genomic library consisted of the following: first, the raw reads of fastq format were filtered through in-house scripts to optimize read number and to reduce artifacts within the data assembly. In this step, clean reads were obtained by removing low-quality reads and reads containing adapter sequence or poly-N from the raw data. In addition, the number of reads from each RAD site were tracked, and RAD sequences above a certain threshold were removed since highly repetitive sequences most likely represent paralogs. Single mismatch derivatives of these highly repetitive RAD sequences were also removed. Finally, low frequency RAD sequences below a threshold (of ≤5) were also removed from further analysis, as the associated paired-end reads would lack sufficient coverage for accurate genotyping (SNP calling).

We processed the clean RADseq reads using the ipyrad 0.9.56 pipeline assembly (Eaton and Overcast, 2020), which is well suited for downstream phylogenetic analyses (Leaché et al., 2015). Since there was no suitable reference genome, we used a *de novo* method to assemble the R1 clean reads. During demultiplexing, restriction sites were trimmed from all reads. All individuals had deep sequencing and high-quality assembly, hence all individuals were retained for downstream analyses. Given the genus *Sinocyclocheilus* is tetraploid (Li and Guo, 2020), we increased the depth of coverage of retained loci to 10x, for accurate genotyping and unbiased heterozygosity. Thus, the maximum number of alleles in the individual consensus sequence was set to 4, to allow more allele combinations within each individual. We set the clustering threshold to 90% similarity and allowed trimmed reads of at least 75bp to proceed in the assembly. In addition, datasets with various minimum numbers of individuals per locus were filtered (96, 102, 108, 114 and 120, corresponding to 80–100% of individual coverage). Other parameters in ipyrad were set to default values.

Demultiplexed, unfiltered reads for the RAD data pertaining to this study can be accessed through the NCBI GenBank Short Read Archive (PRJNA764266).

### 2.3 Phylogenomic analyses

We used the concatenated sequences from all loci of the m114 matrix (8.44% missing data), corresponding to at least 95% individual coverage across loci to infer phylogenetic relationships among all samples. Phylogenomic relationships of all *Sinocyclocheilus* samples were inferred based on the maximum likelihood analysis of a matrix with 1,560,414 characters under a GTRCAT model (Stamatakis, 2014). We also inferred the species tree using the unlinked SNP dataset of the m114 matrix (with a single SNP sampled from each locus) in SVDquartets (Chifman and Kubatko, 2014) implemented in PAUP* 4.0a (Wilgenbusch and Swofford, 2003). SVDquartets estimates the relationships among taxa under the coalescent model by inferring splits among quartets of randomly sampled taxa. All possible taxa quartets (8,214,570 quartets) were evaluated and node support was estimated with 100 bootstrap replicates. The species tree inferred in SVDquartets can assist in identifying incomplete lineage sorting (ILS) or introgression among species in nodes with low support. All phylogenies were visualized using FigTree v1.4.3.

### 2.4 Population cluster analyses

We used a principal component analysis (PCA) to downscale genomic differences among individuals into the first two principal components and visualize the differences between populations. The PCA was based on the .snps.hdf file generated by ipyrad after assembling the data using the ipyrad.pca tool (Eaton and Overcast, 2020). Prior to the analysis, we performed the following treatments: (1) Assigning samples to populations based on the phylogenetic relationships using the imap dictionary. (2) Filtering SNP data using the minmap dictionary to ensure that 50%, 60%, 70%, 80%, 90% of samples have data in each group. (3) Inputting missing data using the “sample” algorithm. Each RAD locus was randomly subsampled to a single SNP to reduce the effect of linkage on the results. After filtering, 3,459, 2,942, 1,993, 987, and 165 SNPs were included in the analysis.

### 2.5 Species delimitation method

We performed Bayes factor species delimitation analyses using BFD* (Leaché et al., 2014). This method computes the marginal likelihood of species trees inferred with the software SNAPP using the Path Sampling approach in BEAST2 (Bouckaert et al., 2014). The method can be used to compare different species delimitation models using genome-wide SNP data in a multispecies coalescent framework. Due to computational limitations, we did not analyze all 120 samples simultaneously. We conducted three independent analyses of species delimitation (Table 1): the first including *S*. cf. *guanyangensis* with the other three closely related taxa in clade A (26 samples), the second including *S*. cf. *longibarbatus* with the other four closely-related taxa in clade C (28 samples), and the third including *S. furcodorsalis* with the other 4 closely-related taxa in clade B (26 samples). The data were filtered to include no missing data among selected samples (Table 1). The parameters for SNAPP were set following Leaché and Bouckaert (2018) for the BFD* analyses (Leaché and Bouckaert, 2018). We conducted a path sampling for a total of 48 steps (MCMC length = 100,000, pre-burnin = 10,000) to calculate the marginal likelihood estimation (MLE) for all models, ranking them by comparing the size of the Bayes Factors across different models.

**Table 1.**
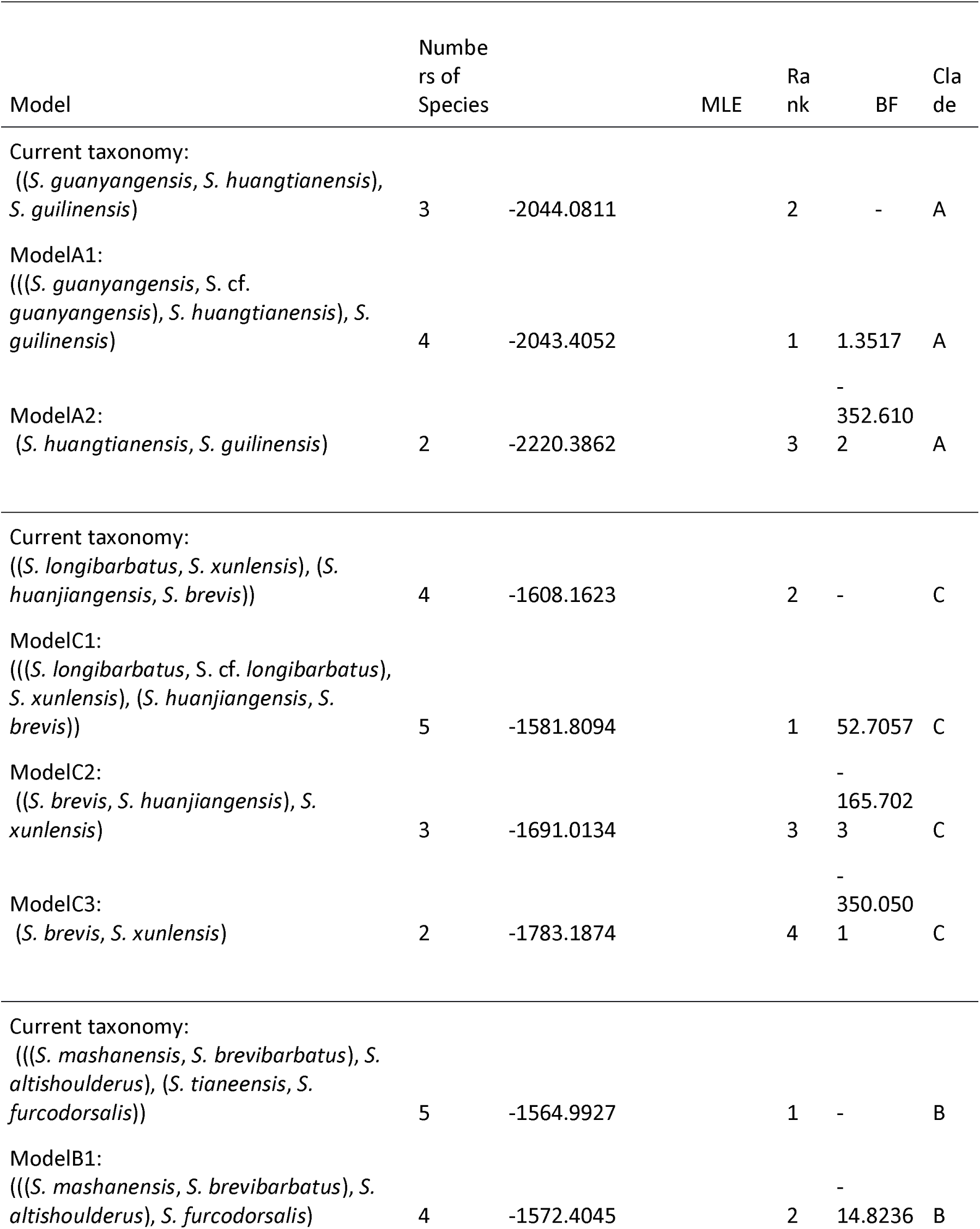
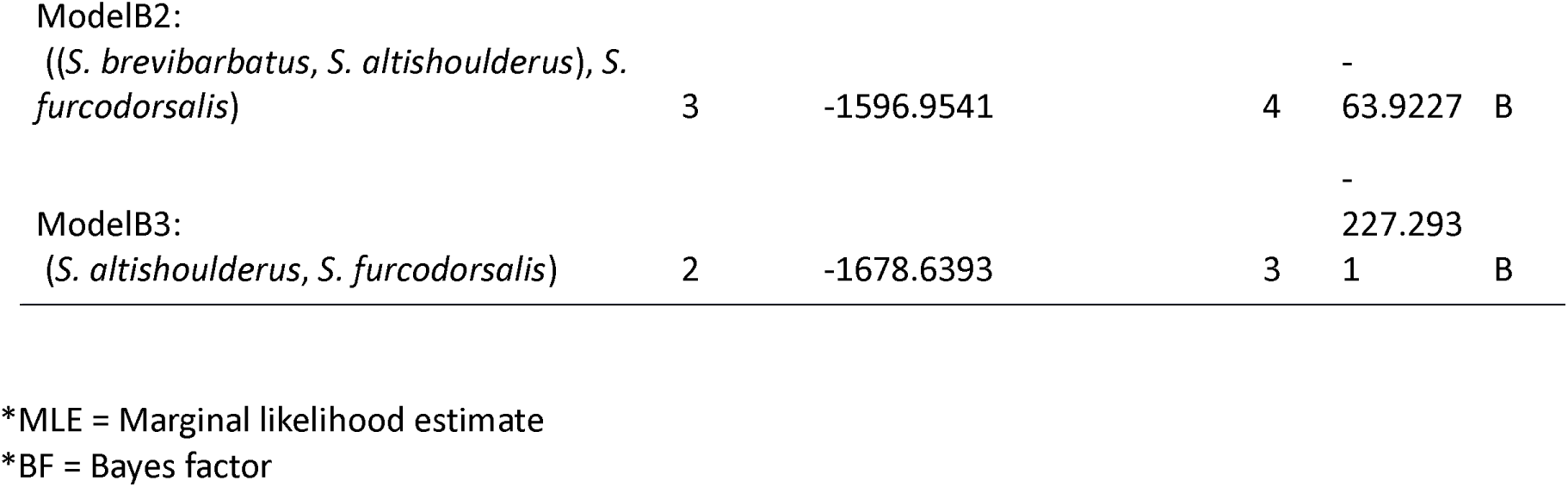
Description of the BFD* species delimitation models for the *Sinocyclocheilus* groups, including the model tested, the number of species, the MLE, the BF between that model and the model of current taxonomy, the model rank and the clade to different species models belong. Since the unidentified species are sister groups with *S. guanyangensis* and *S. longibarbatus*, we herein label them as *S*. cf. *guanyangensis* and *S*. cf. *longibarbatus*.

### 2.6 Divergence time estimation

We inferred divergence times for the *Sinocyclocheilus* using the MSC model in SNAPP, building a species tree of unlinked SNPs following Stange et al (2018), as this algorithm can compensate for ascertainment bias introduced by the exclusion of invariable sites (Bryant et al., 2012; Stange et al., 2018). As running the complete data set was computationally challenging, we selected a single representative per species (21 terminal clades) and reassembled them in ipyrad (2,602 unlinked SNPs for 21 individuals). We then used the Ruby script snapp_prep.rb to prepare the XML input for SNAPP (available at https://github.com/mmatschiner/snapp_prep). For time calibration, we used the time inferred by Liang et al (2011) to the split between *S. donglanensis* and its sister group *S. lingyunensis* at approximately 1.32 Mya (SD:1.02, 95%CI: 0.08-3.91) (Liang et al., 2011). We then ran an independent SNAPP analysis with 20 million MCMC generations, sampling at every 2,000 steps. We checked for stationarity and convergence of chains in TRACER v1.7.1 (ESS>200). A maximum clade credibility tree was generated with TreeAnnotator discarding the first 10% of each MCMC chain as a burn-in. The program FigTree v.1.4.3 was used to visualize the summary tree.

### 2.7 Tests of introgression

To understand the causes of conflicting phylogenetic signals in the species and gene trees of *S. tianlinensis, S. yishanensis*, and *S. macrophthalmus*, we used Treemix from the ipyrad analysis toolkit inferring instances of ancestral gene flow (Pickrell and Pritchard, 2012). Prior to the Treemix analysis, the dataset was processed and filtered in the same way as in the principal component analysis, obtaining 3,459 SNPs. We tested the number of migration edges (m) in the range of 1 to 10, estimating their likelihood score to determine the appropriate value for migration edge in this data. Adding additional admixture edges will always improve the likelihood score, but with diminishing returns as you add additional edges that explain little variation in the data (Kim et al., 2021; Popovic et al., 2020).

## 3. RESULTS

### 3.1. RADseq dataset

The RADseq genomic libraries of the 120 individuals of *Sinocyclocheilus* yielded approximately 9.8 million reads per individual on average, of which 99.91% were retained after the filtering step of the assembly (supplementary table S1). We selected the parameter combination Sino_m114 (supplementary table S2) for phylogenetic analyses as it had the optimal combination of loci-clustering parameters for *Sinocyclocheilus* that maximized the fraction of variable sites that were phylogenetically informative. The data comprised a total of 646,497⍰bp, including 4,378 loci and 67,983 SNPs shared across at least 114 individuals (95% individual coverage at a given locus), of which 61,023 SNPs were parsimony informative for phylogenetic analyses (Sino_m114; supplementary table S2).

### 3.2 Phylogenetic reconstruction of RADseq data

The maximum likelihood analysis of the concatenated RADseq loci in RAxML and the coalescent-based SVDquartets analysis of unlinked SNP data recovered similar phylogenetic relationships among the 19 known *Sinocyclocheilus* species with a high degree of support (> 95) for most nodes (Fig. 1, Fig. 2). As expected, the bootstrap support values at a few nodes of the tree obtained by the coalescent SVDquartets analysis were slightly lower than those obtained by the concatenated analysis (Fig. 1). The two nodes with low bootstrap support in the SVDquartets phylogeny were: the one separating *S. macrophthalmus* and *S. yishanensis* (= 74), and that corresponding to the MRCA of *S. macrophthalmus, S. yishanensis, S. lingyunensis, S. donglanensis* (= 55) (Fig. 2), which suggests some uncertainty in the placement of *S. macrophthalmus* and *S. yishanensis*. Our phylogenetic analyses (Figs. 1-2, A– F) recovered *Sinocyclocheilus* as consisting of six major clades, four of which has been recognized before (Clades A, B, C, and D – Mao et al. 2021). *Sinocyclocheilus tianlinensis*, recognized as a new Clade E, is recovered as the sister species to Clades B, C, D, and F, which is not consistent with the topology of the mtDNA phylogeny (Mao et al., 2021). Likewise, *S. microphthalmus*, recognized as a new Clade F, is now recovered as the sister species to Clades C and D, which is also not congruent with previous phylogenies. Additional incongruences to previous mt-DNA based studies in the Clade C include: *S. longibarbatus* recovered as the sister species to *S. xunlensis*, and *S. yishanensis* as the sister species to *S. macrophthalmus* (Mao et al., 2021). Interestingly, one incongruence also occurred across our trees inferred by different methods: *S. tianlinensis* had a different relationship with a somewhat low bootstrap value in the SNAPP tree (Fig. 3). This suggests conflicting signal, possibly due to introgression. Other than that, relationships across all analyses were congruent for all the remaining species.

**Fig 1.**
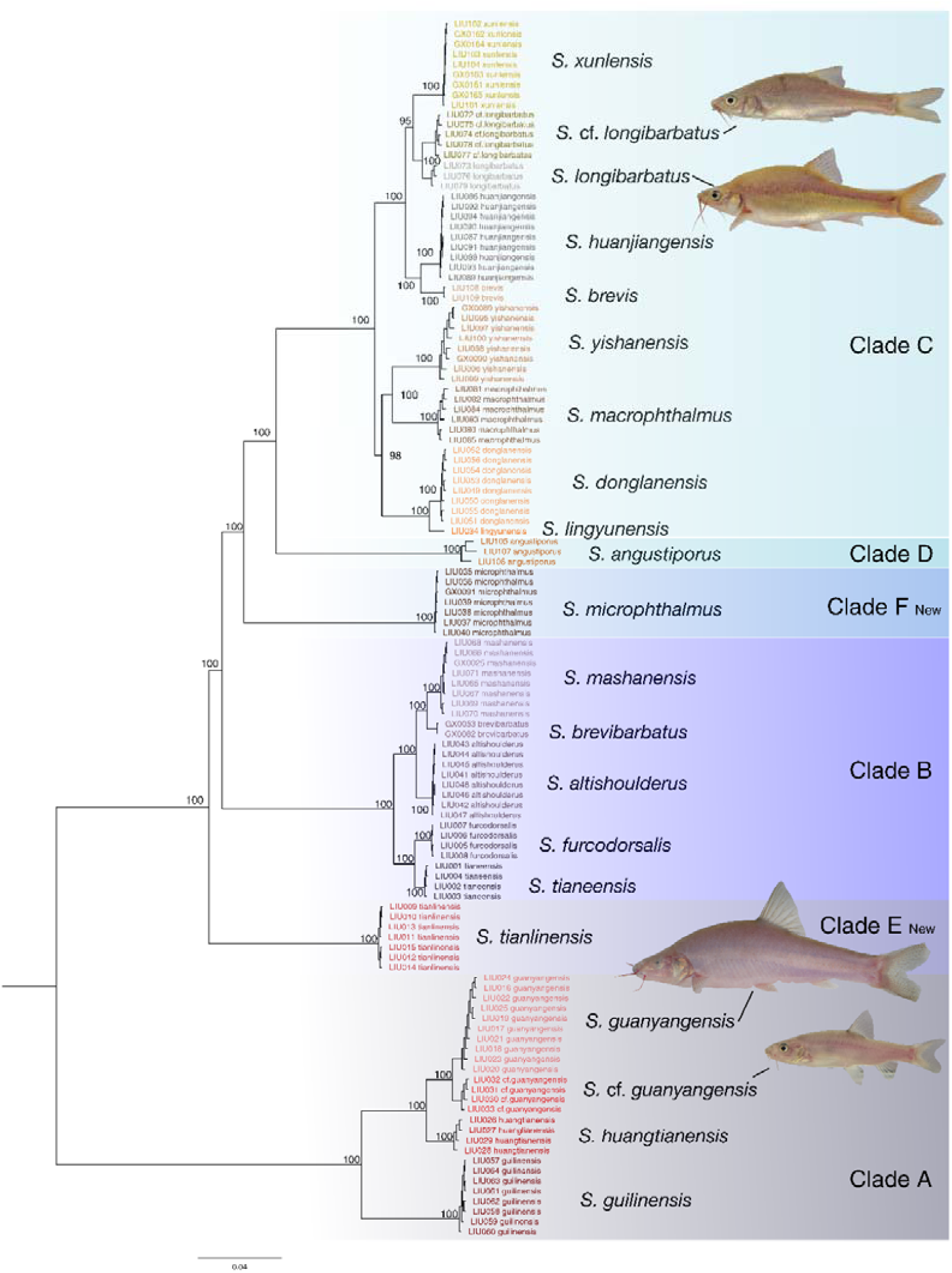
Phylogenomic relationships of the *Sinocyclocheilus* species based on the unpartitioned concatenated maximum likelihood (ML) analysis of a matrix with 646,497 characters using a GTRCAT model of substitution. Bootstrap values are displayed on the nodes.

**Fig 2.**
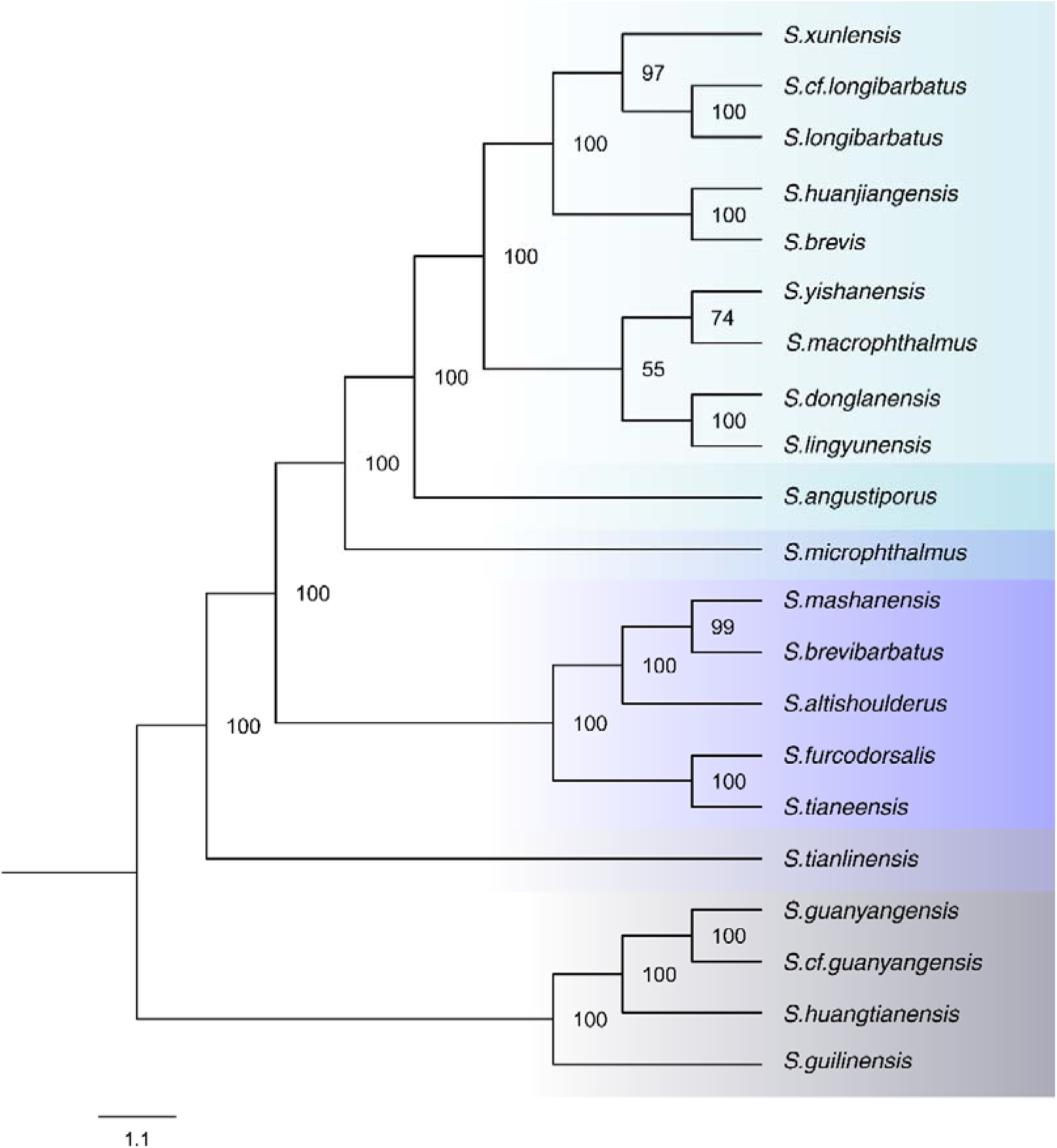
Topology estimated from SVDquartets species tree analysis indicating the phylogenetic relationships of all the 21 *Sinocyclocheilus* species and bloodlines identified in our study based on 4378 unlinked SNPs for 120 individuals.

**Fig 3.**
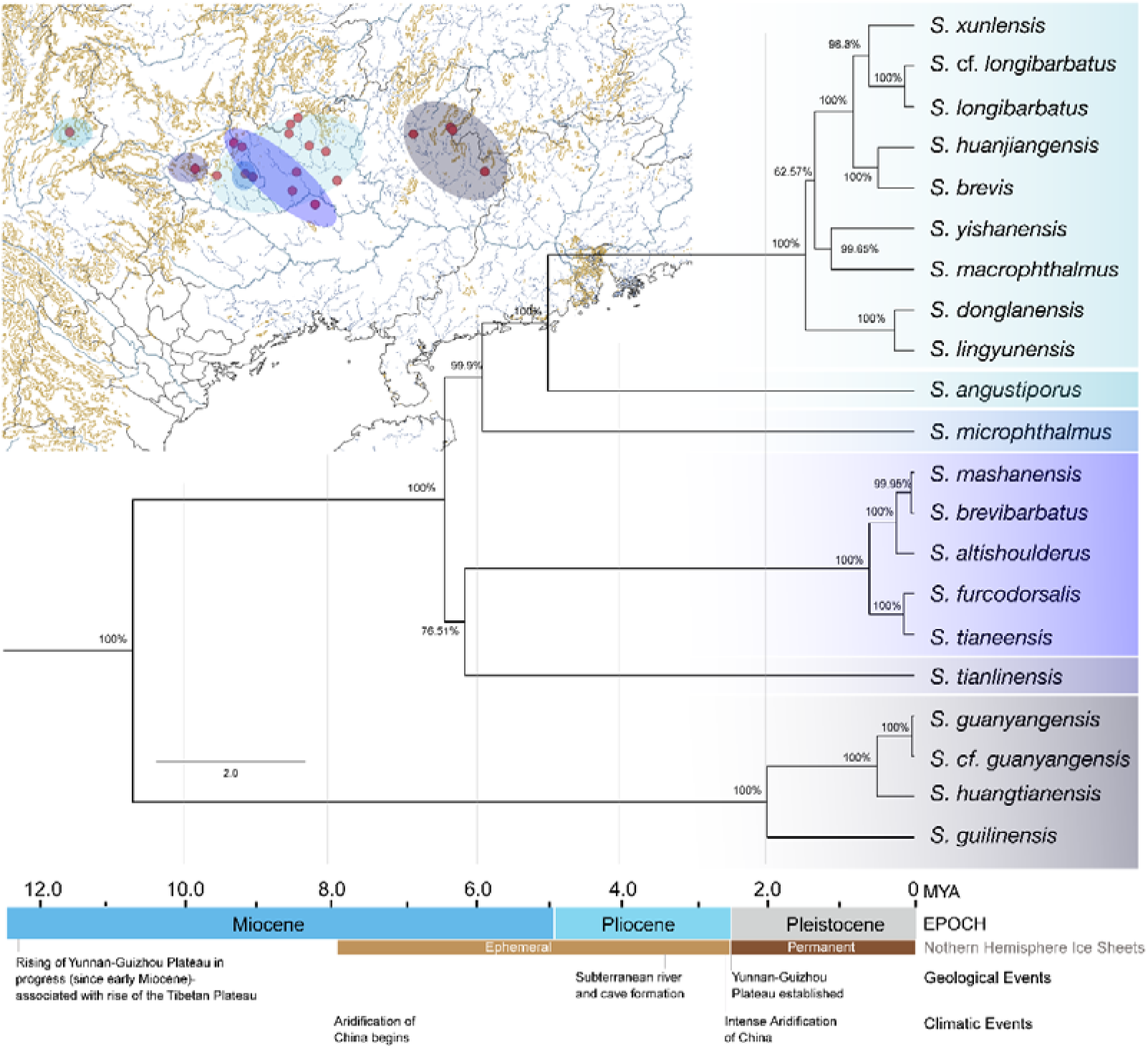
Time-calibrated maximum clade credibility species tree of the all 21 *Sinocyclocheilus* species is inferred by SNAPP. Node support is indicated by the Bayesian posterior probability next to nodes.

### 3.3 Population clustering

We used PCA plots to summarize and visualize in two dimensions the genomic differences across all samples of *Sinocyclocheilus*. The first two principal components explained between 32.2% and 24.9% of the variation using different thresholds for data completeness (50% – 90%) (Fig. 4). The first two axes showed that samples clustered into six subgroups congruent with the major clades identified in our phylogenies.

**Fig 4.**
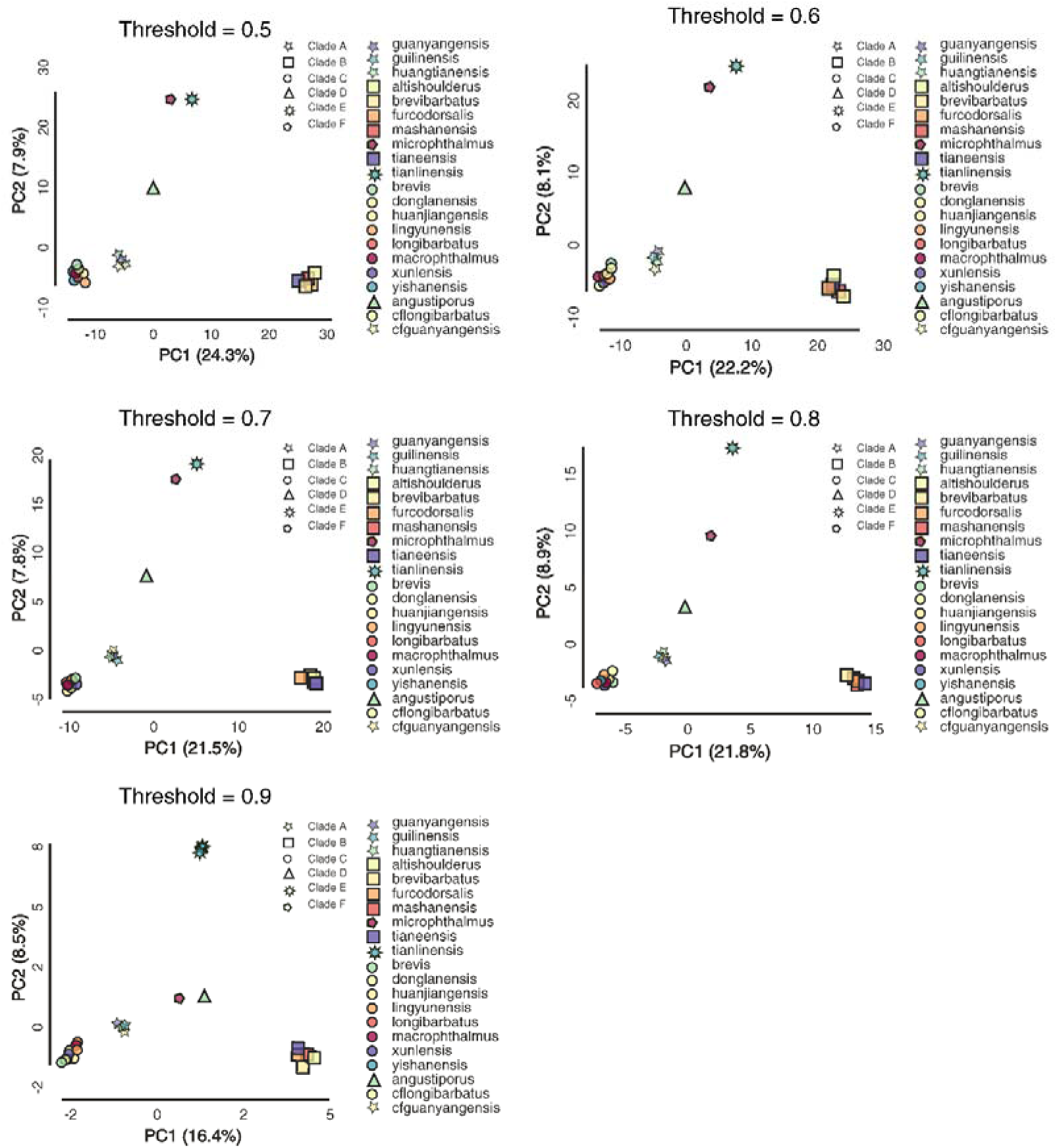
The PCA plots based on the 3459, 2942, 1993, 987, 165 SNP dataset show the population clustering between the 120 samples. The projection of individual samples on the surface is defined by the first two axes of the principal components, the x-axis (PC1 = 24.3%, 22.2%, 21.5%, 21.8%, 16.4%) and the y-axis (PC2 = 7.9%, 8.1%, 7.8%, 8.9%, 8.5%). Different symbols corresponded to the different clades. Different colors corresponded to differentiate the species inside each clade.

### 3.4 Species delimitation

The results for all models tested with BFD* method are summarized in Table 1. Models A1 and C1 support the recognition of *S* .cf. *guanyangensis* (MLE= −2043.41) and *S*. cf. *longibarbatus* (MLE= −1581.81) as separate species. In addition, whether *S. furcodorsalis* and *S. tianeensis* are the same species is also controversial (Liang et al., 2011; Zhao and Zhang, 2009). However, using the BFD* method, the model representing the current taxonomy showed a high Bayes Factor value (MLE= −1564.99), suggesting that *S. furcodorsalis* and *S. tianeensis* are indeed distinct species, supporting the current taxonomy.

### 3.5 Divergence times estimates

Our time-calibrated tree in SNAPP suggests that the crown age for the genus *Sinocyclocheilus* is around 10.5 My old. In an MSC model, *S. tianlinensis* and Clade B show a sister-group relationship (with probability of 0.77). In Clade C, *S. yishanensis* and *S. macrophthalmus* (probability 1.00), *S. lingyunensis* and *S. donglanensis* (probability 0.77) formed sub-clades. (Fig. 3). Compared with the concatenated ML and SVDquartets trees, the SNAPP tree showed a different topology with 5 major clades, including *S. tianlinensis* in Clade B.

### 3.6 Treemix analysis

The likelihoods for different migration boundaries increased steadily as more edges are included in the model (from 0 to 10). However, the likelihood did not improve significantly when the number of migration edges increased above 5. Treemix analyses of m=0–5 suggests the possibility of ancestral gene flow in *S. tianlinensis* and the ancestral species of *S. furcodorsalis* and *S. tianeensis*. Moreover, the possibility of gene flow also exists in the ancestors of *S. yishanensis, S. macrophthalmus* and *S. lingyunensis, S. donglanensis* (Fig. 5).

**Fig 5.**
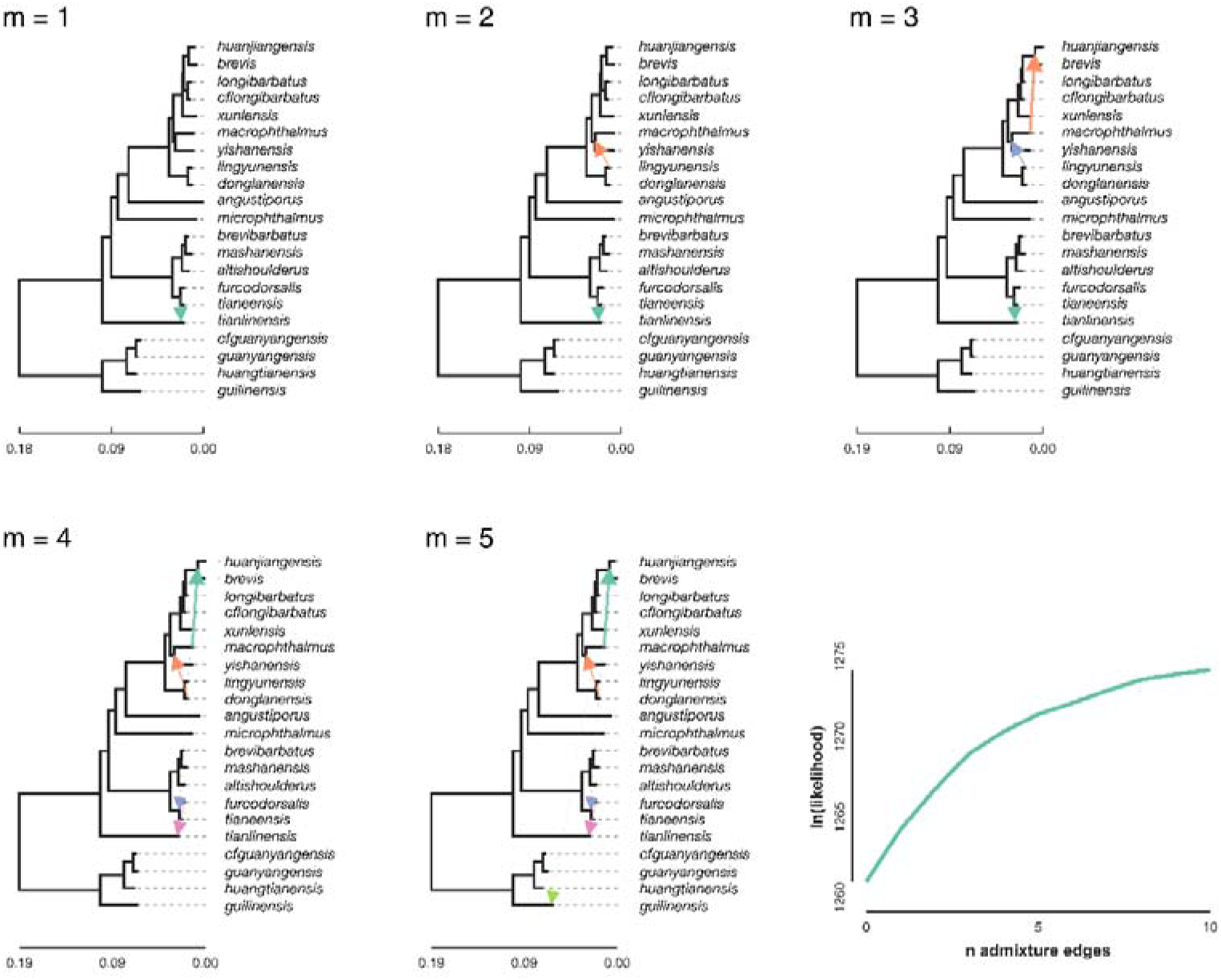
Treemix trees with different admixture edges (m= 1-5) and the plotting of the ln(likelihood) for different values of admixture edges. Arrows indicate when migration occurred in the population tree. The thickness represents the strength of the migration weight.

## 4. DISCUSSION

### 4.1 Phylogeny and cryptic species of *Sinocyclocheilus*

A total of 75 species of *Sinocyclocheilus* fish from China has been described up to now (Jiang et al., 2019), mainly based on morphology and mtDNA data. Nevertheless, some authors have questioned the validity of a few of those species (Liang et al., 2011; Zhao and Zhang, 2009) mainly due to their morphological similarities and incongruences across mt-DNA based phylogenies. For example, the validity of *S. furcodorsalis* and *S. tianeensis* has been questioned by several authors due to their close morphological resemblance (Liang et al., 2011). Such controversies have hindered drawing general conclusions concerning the evolution, ecology, conservation, and biogeography of *Sinocyclocheilus*. Therefore, this genus is in urgent need of a resolved a phylogeny and a clear taxonomy to support robust comparative analyses of the evolution of cave-adaptations in this group.

To this end, we tested the species boundaries in *Sinocyclocheilus* using RAD sequencing-based molecular species delimitation, which had found wide utility in previous evolutionary research (Herrera et al., 2016; Leaché et al., 2014; Rancilhac et al., 2019; Razkin et al., 2016). In addition, *Sinocyclocheilus* had never been subjected to a rigorous molecular species delimitation analysis before. Furthermore, while morphological changes in *Sinocyclocheilus* occur rapidly, molecular divergence, especially in regions under neutral evolution, is assumed to be limited. Therefore, we presumed that mt-DNA based molecular delimitation would not reflect accurate phylogenetic relationships in this group as clearly as the current study, based on a genome-scale sampling that also includes regions under selection. This increased genomic sampling may reflect more accurately the species tree in which *Sinocyclocheilus* evolved their specialized morphology for cave dwelling. Nevertheless, the major conclusions based on the mtDNA phylogeny of Mao et al. (2021) - that blind species independently evolved at least three times - is supported by this RAD-marker based phylogeny as well (Mao et al., 2021).

Trees based on RAXML, SVDquartets and SNAPP were for the most part congruent comprising highly supported nodes with only two nodes on the species trees with low bootstrap support. These species tree methods (i.e. SVDquartets or SNAPP) analyze each SNP separately based on the coalescent theory, so they are expected to show lower bootstrap support whenever there is a conflicting signal among different SNPs in the dataset (Leaché et al., 2015). The tree generated with SNAPP also had a few inconsistencies with the other trees. In the SNAPP tree, *S. tianlinensis* is a sister species to Clade B while *S. donglanensis* and *S. lingyunensis* are sister species to Clade C. However, in the concatenated RAxML tree, *S. tianlinensis* is a sister species to Clades B, C, D, F, while *S. donglanensis, S. lingyunensis* are a sister clade to *S. macrophthalmus* and *S. yishanensis* in Clade C. We assume that this conflicting phylogenetic signal is either due to ancestral gene flow, introgression or incomplete lineage sorting. Treemix analysis suggests introgressive gene flow.

Previous evolutionary studies based on mitochondrial genes have shown that *Sinocyclocheilus* is divided into four major clades (Liang et al., 2011; Ma et al., 2019; Romero et al., 2009; Zhao and Zhang, 2009). However, the phylogenetic relationship obtained with RAD-seq analysis in the present study suggests that this group can be further divided into two additional branches, comprising six major clades (Fig. 1). The principal component analysis further supports this division by clustering the samples into six clusters which correspond to the six clades (Fig. 4). Previous mt-DNA based phylogenies (Mao et al., 2021) recognized *S. tianlinensis* and *S. microphthalmus* as species within clade B, but our phylogenomic results indicate that those two species actually belong to two independent clades due to their deep divergences in Clades B and C respectively.

Our results also support that *S. guanyangensis* and *S*. cf. *guanyangensis*, which resemble each other morphologically, are in fact two distinct species, the latter being a putative new species. The same is true for *S. longibarbatus* and *S*. cf. *longibarbatus* (a putative new species) as well. Field sampling indicated that these species pairs occur in different caves, but in close proximity to each other (see map in Fig. S1). Based on morphological and molecular data, it appears that different species of cave fish have repeatedly invaded different cave waters and acquired their own independent troglomorphic characteristics. This may be the case for these species as well. Geographical isolation followed by strong genetic drift among populations seems to be the main mechanism for the rapid formation of species in the genus *Sinocyclocheilus* (Ma et al., 2019; Yang et al., 2021; Zhao and Zhang, 2009). However, especially in small, isolated populations such as *S*. cf. *guanyangensis* and *S*. cf. *longibarbatus*, genetic differentiation and species formation may occur even more rapidly due to the lack of gene flow among isolated populations. This mechanism, occurring under strong selective pressure associated with cave dwelling, may accelerate and amplify the rapid evolution of adaptive traits in other species of *Sinocyclocheilus* as well, resulting in a particularly rich species diversity with frequent evolution to cave-adapted morphologies.

Our genome-wide SNP analyses identified two putative new species of *Sinocyclocheilus* out of the 120 individuals sampled across 21 localities. Considering that our study did not include samples for all 75 species described so far, we suspect that cryptic diversity in the group could be potentially higher, i.e., the species diversity of the *Sinocyclocheilus* complex may be vastly underestimated.

### 4.2 Divergence time estimation and gene flow

The age of the most recent common ancestor (MRCA) of *Sinocyclocheilus* fish, inferred by our RADseq data, was estimated to be around 10.5 Mya, similar to that inferred by previous mtDNA-based studies (Li et al., 2008; Liang et al., 2011; Mao et al., 2021). We also found that most divergence events occurred relatively recent in the history of the group (in the last 2 Mya) (Fig. 3). The *Sinocyclocheilus* MRCA possibly lived on the Yunnan-Guizhou Plateau during the late Tertiary. During the Quaternary, the Qinghai-Tibetan Plateau underwent an abrupt upturn, with major changes in the geological environment and a dramatic transformation of the Yunnan-Guizhou Plateau (Li and Fang, 1999; Shi et al., 1999). At the same time, global temperature began to fall and Northern Hemisphere ice caps grew (Hewitt, 2000; Ma et al., 2019; Svendsen et al., 2004). The MRCA of the *Sinocyclocheilus* radiation may have adapted to live in underground caves due to dramatic changes in the environment. The relatively late tectonic uplift of the Tibetan Plateau 3.6 Mya may have affected population dynamics of *Sinocyclocheilus* as well. For example, three species (*Sinocyclocheilus grahami, S. rhinocerous*, and *S. anshuiensis*) experienced two episodes of population declines during the two intense uplift phases (Qingzang movement: 3.6 Mya∼1.7Mya, Kunhuang movement: 1.1Mya∼0.6Mya) of the third tectonic uplift of the Qinghai-Tibet Plateau (Yang et al., 2016). Changes in population size and gene flow may be related to enhanced Asian monsoon and precipitation, with large-scale glacial activity resulting from these two phases considerably affecting diversification in this group. Therefore, the intense late tectonic activity in the Qinghai-Tibet Plateau, combined with environmental changes, could have promoted the rate of diversification in these cavefish during the last 2 My.

SNAPP is a multispecies coalescent method that assumes no gene flow among species. This may lead to inaccurate inference of topology in the presence of gene flow and may explain the incongruences among the SNAPP tree and the other trees. In the Treemix analysis, we found possible ancestral gene flow between some species of *Sinocyclocheilus*, which is likely responsible for the inconsistent phylogenetic signals described previously (Fig. 5). Subterranean river-capture events may have influenced this phenomenon. The geographical locations of the species showing possible ancestral genetic admixture are in close proximity and were possibly connected through underground networks, especially during the Pleistocene. With the upliftment of the Tibetan plateau during the early Pleistocene, many cave systems within the basin, which were interconnected previously, would have become completely or periodically isolated, preventing gene flow among populations in close proximity, giving rise to the patterns among sister taxa that we see today. Interestingly, gene flow is more prominent around the karstic northwestern regions of the Guangxi plains, especially in Clades B and C. Therefore, we can assume that the deeper caves and the subterranean river systems associated with the Guangxi karst region (Zhao and Zhang, 2009) were periodically interconnected, allowing limited gene flow. The deeper phylogenetic relationship of *S. tianlinensis* and the species in Clade B, suggests that they diverged during the Miocene. Furthermore, introgression is not apparent among the species from the hilly terrains of Yunnan plateau, suggesting that they inhabit isolated subterranean systems in these hills.

## 5. CONCLUDING REMARKS

In summary, our study showed that: 1) instead of the 4 major clades of *Sinocyclocheilus* previously recognized by phylogenetic relationships, the genus can be now categorized into 6 major clades; 2) the MRCA of *Sinocyclocheilus* appeared around 10.5 Mya coinciding with cave formation due to upliftment and dry conditions associated with the aridification of China during the late Miocene and the Pliocene; 3) the BFD* analyses support the hypothesis that the two cryptic species found in this study (*S*. cf. *longibarbatus* and *S*. cf. *guanyangensis*) and the morphologically similar *S. tianeensis* and *S. furcodorsalis* are distinct species. We further draw attention to the fact that the diversity of the *Sinocyclocheilus* species may be substantially underestimated due to their cryptic nature. Future studies using novel genomic techniques (such as RADseq or whole genome sequencing) could potentially unravel the true diversity of this remarkable radiation.

## Abbreviations

RADseq: Restriction site-associated DNA sequencing
mt-DNA: Mitochondrial DNA
BFD: Bayes factor delimitation
Mya: Million years ago
ILS: Incomplete lineage sorting
PCA: Principal component analysis
SNP: Single-nucleotide polymorphism
MSC: Multispecies coalescent
MLE: Marginal likelihood estimate
ML: Maximum Likelihood

## Acknowledgements

We thank Rohan Pethiyagoda (Australian Museum) and Mario van Gastel for suggestions to improve the paper. We thank the following for assistance in the field: Shipeng Zhou, Bing Chen, Dan Sun, Jayampathi Herath and Amrapali Rajput.

## Authors’ contributions

MM, MMV, MRP, TRM, YWL, conceptualized the research and designed the methodology. YWL, CHF, MTR, MM, GE, JY conducted fieldwork and curated the data. TRM, MMV, YWL, MRP, GE, carried out formal analysis. TRM, MMV, MM, MRP, GE, YWL, wrote the original draft. MM, MMV, MRP, and JY supervised MTR & YWL. MM and JY acquired funding. TMR, GE and YWL made figures. All authors reviewed and edited the draft. All authors read and approved the final manuscript.

## Availability of data and materials

All data generated or analyzed during this study are included as supplementary information and the genetic data (Genbank) can be accessed upon acceptance of the paper. Please see the materials and methods section for the SRA project name for RAD data.

## Ethics approval and permission

Methods of sampling approval by the Ethics Committee of Guangxi University. Field sampling approval through the Guangxi Provincial Government.

## Consent for publication

Not applicable.

## SUPPORTING INFORMATION

**Fig S1.**
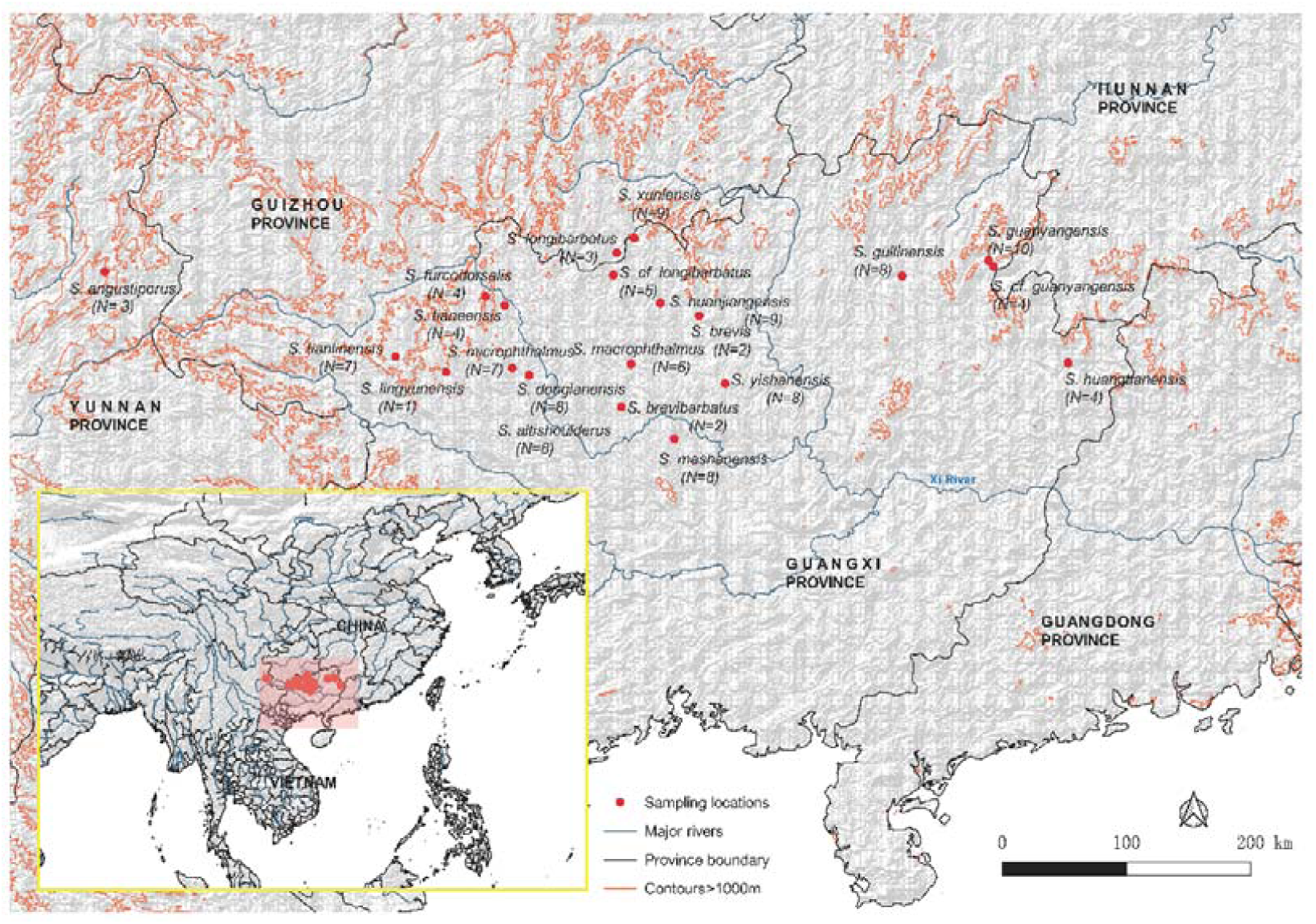
Map of the sampling localities of *Sinocyclocheilus* species. *S.donglanensis* and *S.altishoulderus* are caught in the same cave.

**Table S1.**
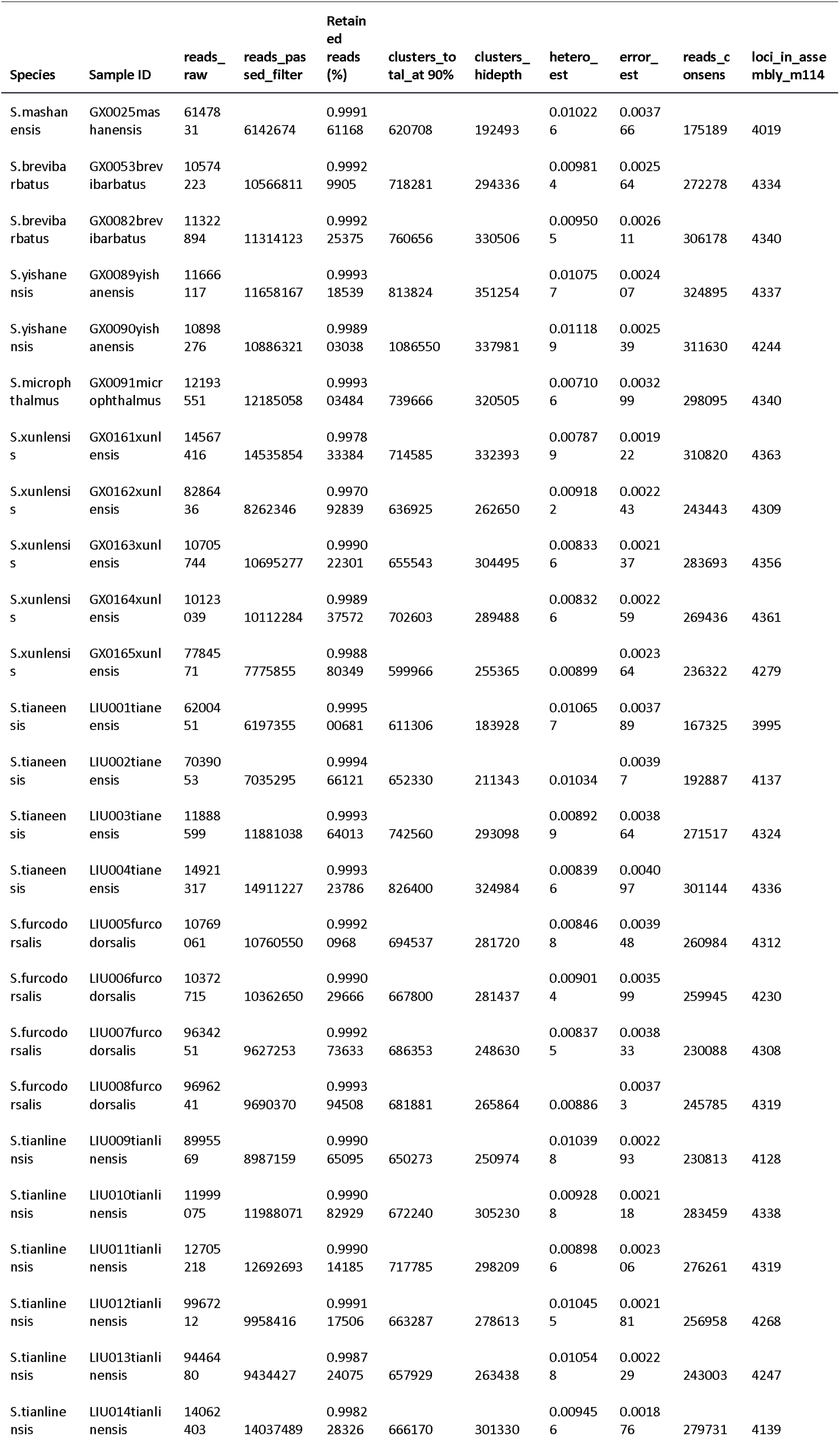
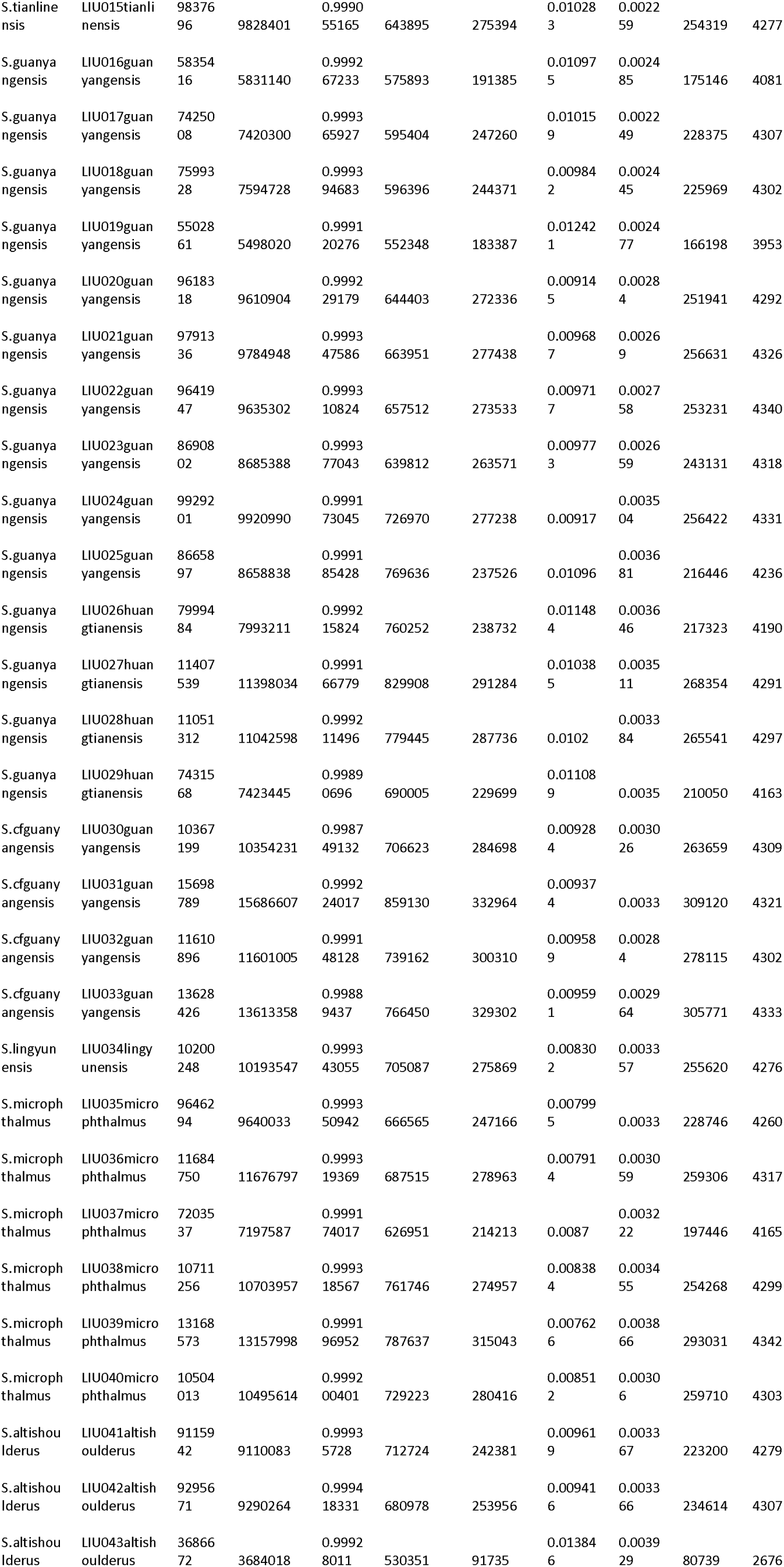
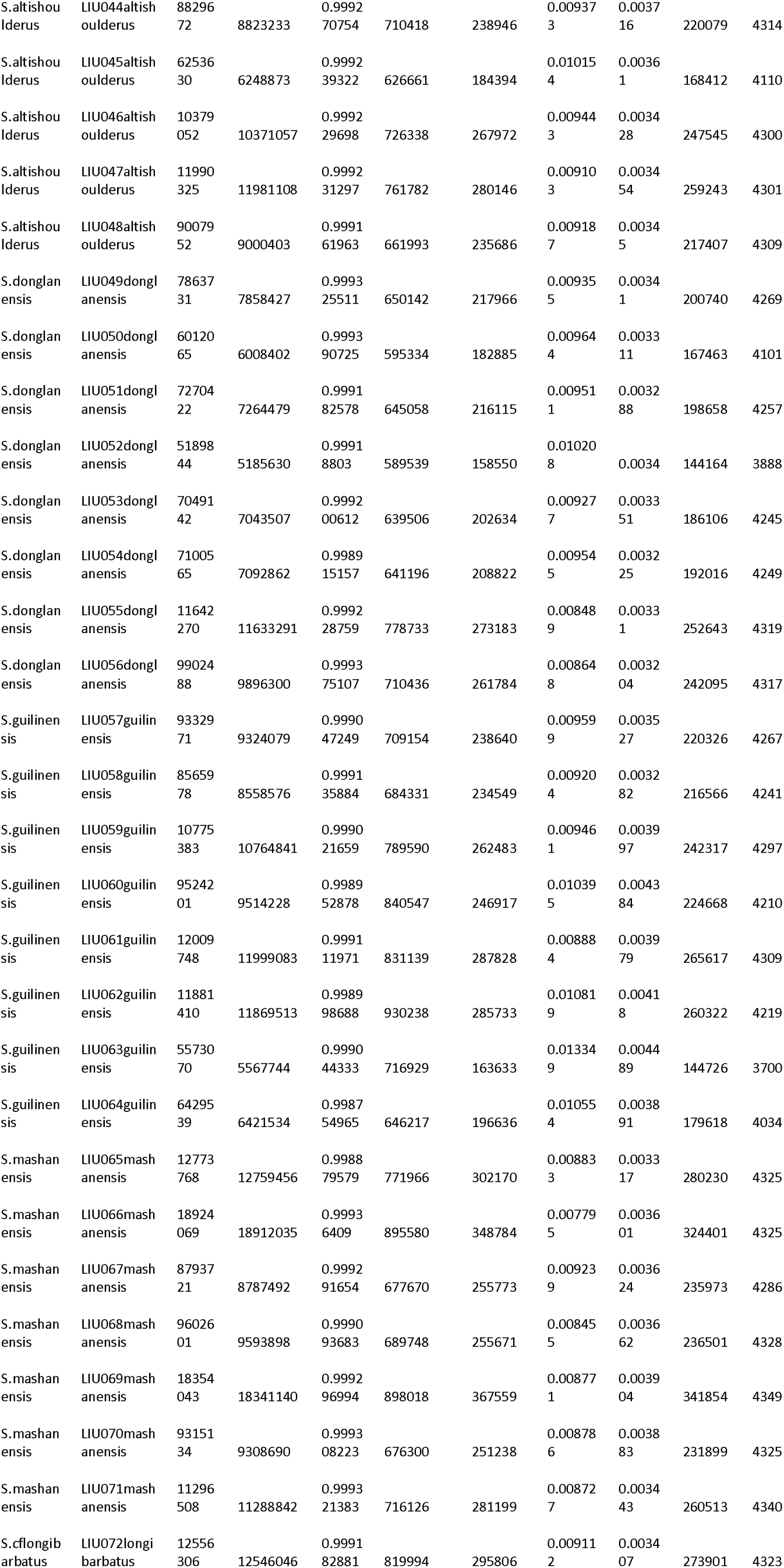
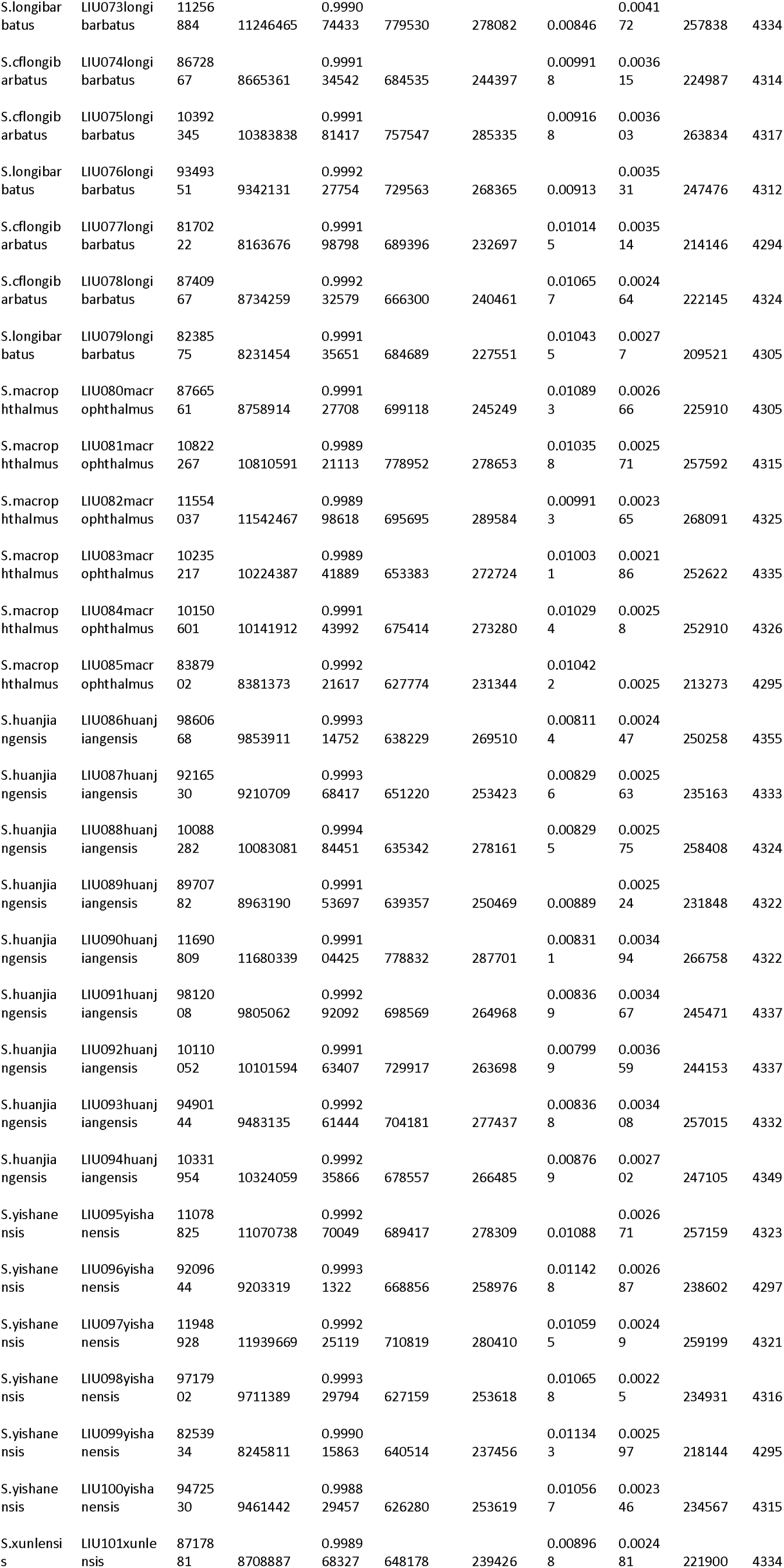
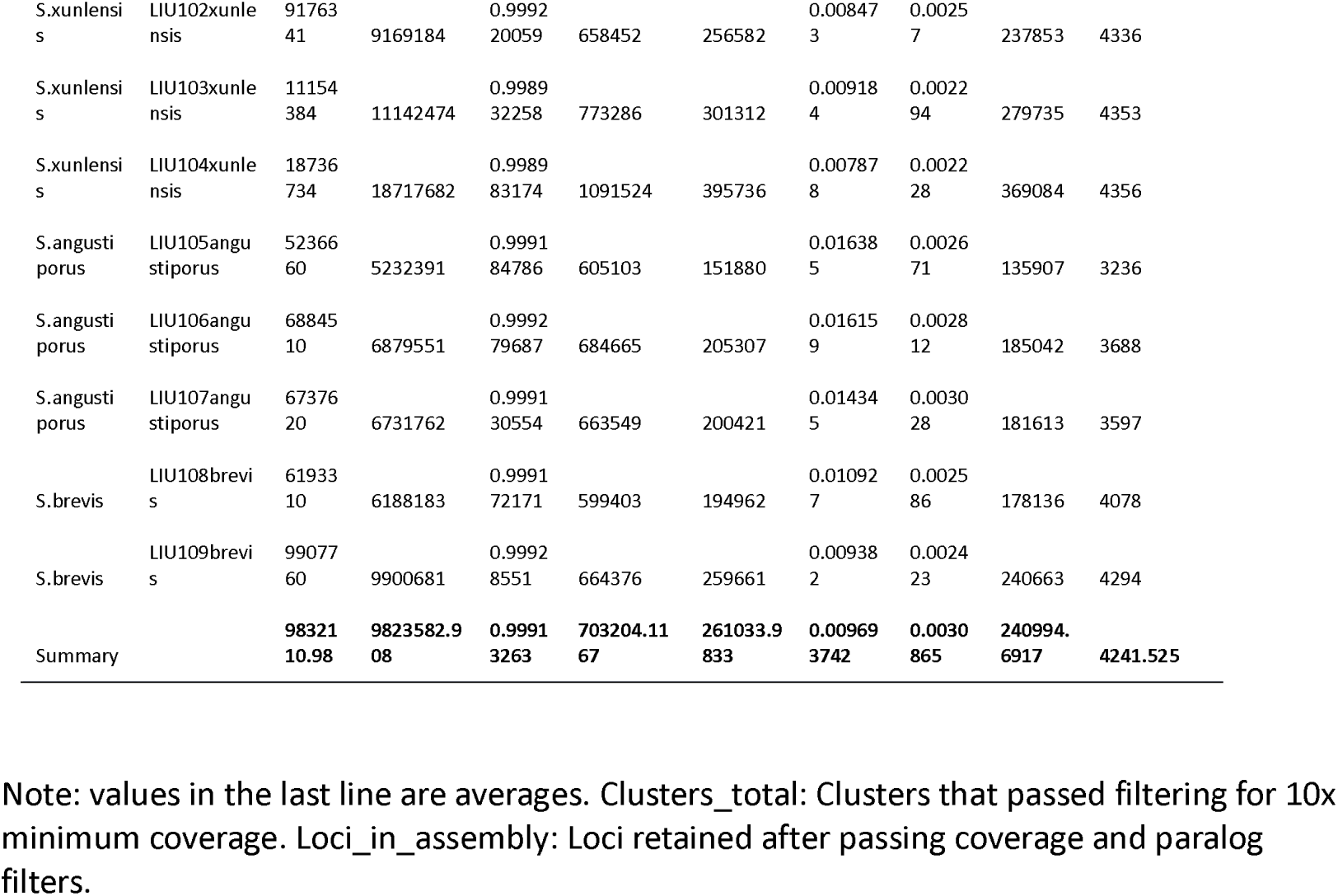
Summary of the RAD data.

**Table S2.** Sequence information with different parameters.

| Matrix | Unlinked SNPs | Consensus sequences (bp) | VAR | PIS | Missing (%) | PIS/VAR(%) |
| --- | --- | --- | --- | --- | --- | --- |
| Sino_m96 | 23,869 | 3,534,842 | 380,519 | 336,928 | 13.46 | 88.54 |
| Sino_m102 | 17,187 | 2,544,628 | 272,838 | 242,335 | 10.89 | 88.85 |
| Sino_m108 | 10,549 | 1,560,414 | 166,407 | 148,253 | 8.21 | 89.09 |
| **Sino_m114** | **4,378** | **646,497** | **67,983** | **61,023** | **5.25** | **89.76** |
| Sino_m120 | 151 | 22,274 | 2,199 | 1,989 | 2.29 | 90.45 |

Note: Sequence information in the RAD data matrices (n=120) generated with different parameters of minimum number of individuals per locus (m) values. The data matrices shown in bold type were used for analyses.

